# A user-friendly goniometer-compatible fixed-target platform for macromolecular crystallography at synchrotrons

**DOI:** 10.1101/2025.09.04.674375

**Authors:** Swagatha Ghosh, Analia Banacore, Per Norder, Monika Bjelcic, Arpitha Kabbinale, Padmini Nileshwar, Gabrielle Wehlander, Daniele de Sanctis, Shibom Basu, Julien Orlans, Adams Vallejos, Leonard M. G. Chavas, Richard Neutze, Gisela Brändén

**Affiliations:** Department of Chemistry and Molecular Biology, Gothenburg University, Sweden; Department of Applied Physics, Graduate School of Engineering, Nagoya University, Japan; BioMAX Beamline, MAXIV laboratory, Lund University, Sweden; European Synchrotron Radiation Laboratory, Grenoble, France; European Molecular Biology Laboratory, Grenoble, France; Nagoya University Synchrotron Radiation Research Center, Nagoya University, Japan

## Abstract

Fixed-target platforms provide convenient support for microcrystals during serial X-ray crystallography studies using synchrotron radiation. Here, we describe a simple, user-friendly 3D-printed support where the crystals are sandwiched between two layers of thin X-ray transparent membrane resulting in very low scattering background. The platform is compatible with magnetic mounting onto the standard goniometer of macromolecular crystallography beamlines. Our design utilizes a 96-well frame that facilitates hanging-drop experiments directly on the membrane using conventional crystallization plates, thereby eliminating multiple pipetting and crystal handling steps. Crystals can be enclosed into a sandwich and packed into ‘cassettes’, preventing the risk of the sample drying out during room-temperature transportation to synchrotron sources. The versatility of the platform is demonstrated by five structures solved using different crystallization- and data-collection strategies. Single crystal rotational-crystallography at both room- and cryogenic-temperatures using large crystals of lysozyme is shown. On-chip microcrystallization is illustrated by use of a photosynthetic reaction center as an example. Finally, serial crystallography data collection at room-temperature from microcrystals of photosynthetic reaction center as well as cytochrome *c* oxidase crystallized in lipidic cubic phase is presented.

## 1. Introduction

X-ray crystallography remains a widely used and well-established structural biology technique for obtaining high-resolution, atomic-level structures of macromolecules (Brito & Archer, 2020). A majority of the protein structures deposited to the protein databank (PDB) emerge from X-ray crystallography owing to its wide-spread use and the efficiency of the method where several of the steps can be automated, including the use of crystallization robotics as well as automation of data collection and data-processing. Moreover, as the crystals are kept frozen, they can easily be shipped to a synchrotron facility allowing remote data collection. However, conventional X-ray crystallography typically requires protein crystals that are at least tens of micrometers in size, manual crystal handling and cryogenic data-collection conditions to limit radiation damage.

Serial X-ray crystallography (SX) was initially developed at X-ray free-electron laser facilities (XFELs) (Chapman *et al*., 2011)and is now established also at synchrotron radiation facilities (Gati *et al*., 2014). Serial data-collection methods allow high-resolution room-temperature structural information to be obtained and has expanded the capabilities for time-resolved structural studies to track protein dynamics (Pearson & Mehrabi, 2020, Branden & Neutze, 2021). In SX, X-ray diffraction images are collected from a large number of randomly oriented small crystals where each crystal that is exposed to the X-ray beam gives rise to one diffraction image. Typically, thousands of images are then indexed, integrated and merged to obtain a complete dataset. As each crystal is exposed to a single short X-ray pulse, there is potential to minimize radiation damage. Moreover, combining data from numerous small crystals allows structure determination of micro- and nanocrystals, may aid in the study of challenging targets like membrane proteins (Weierstall, 2014). As a hybrid alternative using lower-intensity synchrotron beams, narrow wedges of data can be collected from a smaller set of crystals at cryo- or room-temperature (Zander *et al*., 2015). The recent development of purpose-built SX synchrotron beamlines with micro-focus X-ray beams and fast detectors is now allowing many more users to access the method.

SX sample delivery is dependent on continuously replacing the crystals and can be broadly categorized into two main branches: moving-target and fixed-target approaches. Moving targets include the injection methods that involve a continuous flow of microcrystals across the X-ray beam e.g. by use of a liquid- or high-viscosity jet, a capillary, or a microfluidic chip (Weierstall, 2014, Ghosh *et al*., 2023, Monteiro *et al*., 2020). Diffraction images are collected as crystals flow across the beam one after the other (Doppler *et al*., 2022). This approach enables continuous sample replenishment, allowing data collection from a large number of crystals without manual intervention and at a high repetition rate. It is often the method of choice for collecting time-resolved X-ray diffraction data, where a reaction is initiated in the sample at a selected time-point before the X-ray probe. The majority of time-resolved SX studies to date have been of light-activated proteins where a laser trigger is used to start the reaction (Branden & Neutze, 2021, Khusainov *et al*., 2024). A more universal triggering method is by mixing the crystalline sample with a substrate, although the time-resolution achieved can be limiting (Monteiro *et al*., 2020). A severe limitation of the injection methods is that the sample consumption can be very high, especially for the collection of time-resolved data (Lyubimov *et al*., 2015).

An attractive alternative to the injection devices for sample delivery is the use of so-called fixed-targets, also called “chips” (Roedig *et al*., 2015). Fixed-target approaches involve distributing the crystals on a solid support such as a membrane or mesh that is then mounted on a specialized sample holder and raster-scanned across the X-ray beam. Their main advantages are that they allow minimal sample consumption and that the complexity of the experimental setup may be reduced. Depending on the choice of device, crystals can be either in solution phase or in a high-viscosity media such as lipidic cubic phase (LCP) (Berntsen *et al*., 2020). Drawbacks include potential increased background scattering from the support and, in some cases, the need for precise scanning mechanisms at the beamline. In addition, sample handling during the loading of the chip may cause physical damage to the crystals (Martiel *et al*., 2019). Finally, the use of fixed-targets can be limiting for time-resolved studies, although there are set-ups that work very well in combination with both light-triggering (Caramello & Royant, 2024, Schulz *et al*., 2022) and mixing (Mehrabi *et al*., 2020). Two conceptually different variants of fixed-target devices have been developed: aperture aligned or sequentially exposed/directed raster (Carrillo *et al*., 2023). The aperture-aligned approach relies on a precise location of the crystals on the chip (Owen *et al*., 2023). This can be achieved through the use of micro-patterned chips where the crystals arrange themselves in the wells and excess solution is blotted away (Roedig *et al*., 2016). One of the earlier examples is the Roadrunner (Roedig *et al*., 2017) which utilizes a dedicated sample stage and a humidity chamber to keep the crystals from drying out. The well spacing and pore size is chosen to match the crystal size but using very small or thin crystals is problematic as they may escape through the pores. A more recent development is the micro-structured polymer (MISP) chip that allows a cheaper and sturdier alternative (Carrillo *et al*., 2023). The sequentially exposed variant of fixed-targets does not depend on placing the crystals in specific locations, instead they are randomly distributed on the support. These are typically simple designs where the protein microcrystals are sandwiched between two membranes and data are collected by raster-scanning over a pre-defined grid on the chip (Owen *et al*., 2017). An example is the hermetically sealed sheet-on-sheet (SOS) chip that can be used at both XFELs and synchrotron beamlines (Doak *et al*., 2018). To limit the crystals from moving around on the chip during data collection, a variant with 10–15 %(*w*/*v*) viscous gelatin and 1–4 %(*w*/*v*) agarose gel on the membrane was developed (Lee, 2020). There are also tools available that allow data collection under anaerobic conditions (Rabe *et al*., 2020, Bjelcic *et al*., 2023). Most purpose-built SX synchrotron beamlines, including MicroMAX at MAX IV, ID29 of the ESRF (Orlans *et al*., 2025), T-REXX of Petra III and I24 of Diamond, offer access to various fixed-target devices, some of which are summarized in Table 1 with the characteristics of each device indicated.

One of the previous major limitations with the SX method was the challenge of producing suitable and sufficient amounts of microcrystals. Batch crystallization with seeding has proven to be an efficient method for proteins in solution (Dods *et al*., 2017, Dunge *et al*., 2024, Shoeman *et al*., 2023) and approaches for large-scale LCP crystallization of membrane proteins have been presented (Andersson *et al*., 2019). A risk associated with crystallization is that of damaging crystals during handling. Therefore, it may be advantageous to integrate crystallization with data collection, for example through *in situ* crystallization where the crystals are directly grown on the support that is used for data collection (Foos *et al*., 2024). Some crystallization plates allow direct data collection at room-temperature, where typically a wedge of data is obtained through a small rotation of the plate in the beam (Axford *et al*., 2012), and specialized plates enable the collection of diffraction data also under cryo-conditions (Broecker *et al*., 2018). *In situ* crystallization to screen for suitable crystallization conditions has also been achieved in microfluidics chips (Sui, 2017). That said, *in situ* crystallization is not routinely performed in combination with SX fixed-target devices.

Despite the rapid developments over the last decade, many challenges for making the method easily accessible to new users remain. There are issues associated with transportation of crystals at room-temperature which means that crystals in many cases have to be prepared on-site at the radiation facilities. Data collection devices need to be more user-friendly and affordable. Finally, SX data collection would benefit tremendously from the use of robotics to automatically mount the fixed targets at the beamline. In this work, we present a flexible and easy to use 3D printed fixed-target platform. It allows on-chip crystallization, can be transported safely, is pre-assembled to aid sample loading, is compatible with data collection at all common synchrotron beamlines without specialized hardware, and gives high-quality SX data at room-temperature as well as under cryo-conditions. The versatility of the device is showcased by presenting structural data collected at three different synchrotron facilities of three protein systems including a membrane protein crystallized in LCP.

**Table 1.** Examples of fixed-target devices.

| base | *in situ* | disposable | collection strategy | RT | cryo-temp | sample-loading system | reference |
| --- | --- | --- | --- | --- | --- | --- | --- |
| SPINE | no | no | raster | yes | yes | yes | (Illava *et al.*, 2021) |
| customized | yes | no | raster | yes | no | no | (Ren *et al.*, 2020) |
| customized | yes | unknown | directed | yes | no | yes | (Mueller *et al.*, 2015) |
| SPINE | yes | unknown | raster | yes | yes | no | (Lieske *et al.*, 2019) |
| customized | no | yes | raster | yes | unknown | yes | (Lee, 2020) |
| customized | no | partially | directed | yes | unknown | yes | (Carrillo *et al.*, 2023) |
| customized | no | partially | directed | yes | unknown | yes | (Wierman *et al.*, 2019) |
| customized | no | yes | raster | yes | yes | no | (Doak *et al.*, 2018) |
| customized | no | partially | raster | yes | yes | no | (Rabe *et al.*, 2020) |
| customized | no | no | raster | yes | yes | no | (Bjelcic *et al.*, 2023) |
| SPINE | no | no | raster | yes | yes | no | (Park *et al.*, 2020) |
| customized | no | no | directed | yes | yes | yes | (Mehrabi *et al.*, 2020) |
| SPINE | no | yes | raster | yes | yes | no | (Roedig *et al.*, 2015) |
| SPINE | yes | yes | raster | yes | yes | no | this work |

“Base” refers to whether the fixed-target requires a customized base or if it is compatible with the standard SPINE base for data collection (Beteva *et al*., 2006). “*In situ*” refers to whether *in-situ* crystallization is possible and “disposable” whether the device is disposable or reusable. “Collection strategy” describes if X-ray data are collected using a raster scan or if the crystals are in pre-defined positions (“directed”). “RT” and “Cryo-temp” refers to that the data can be collected at room-temperature and cryo-temperature, respectively, and “sample loading system” whether a custom-made system is needed to load the sample onto the chip. Mehrabi *et. al.* describe the so-called HARE chip and Roedig *et. al*. describe a silicon chip commercially available from Silson. The fixed-target device described in this study is highlighted in green.

## 2. Materials and methods

### 2.1 Design of a chip-based framework for hanging drop crystallization, crystal transportation and data collection

The chip-based platform described here consists of three parts: 1) a framework consisting of ‘chips’ for hanging-drop crystallization and/or encapsulation of crystals, 2) a compact device for crystal transportation, and 3) a goniometer-compatible holder for X-ray diffraction data collection. The models for different parts of the platform were prepared using computer-aided design (CAD) technology with AutoCAD (Autodesk 2020) and printed in our in-house 3D printers Asiga MAX/MAX UV using plastic resin (PlasGRAYV2) and/or FELIX Pro 3 printer (with polylactic acid, PLA filaments). Firstly, 3D-models of a solid frame containing a circular disk (or hole) that could fit onto a reservoir of a standard 24-well (Hampton Research) or a 96-well (Greiner and TPP) hanging drop crystallization plate are printed. Each disk is then layered with an X-ray transparent membrane (Mylar or Kapton) of variable thickness (3-12 microns) using an adhesive. Double-sided adhesive films accurately fitting the size of the disks are carved on a Silhouette cameo-4 cutting machine. The 3D-printed frame with disks, double-sided adhesive films and an X-ray transparent membrane is assembled as shown in Figure 1A into an entity called a ‘chip’. For ease of assembly, the 96-well design makes up an array of chips that could fit over a standard crystallization plate. Post-assembly, the frames are wiped with ethanol and dried before being used for crystallization or data collection. Crystalline samples are mounted by closing two chips to form a sandwich (Figures 1B, 2C) with a total spacer thickness of ∼75 micrometers that is intercepted by the X-rays. For storage and transportation, different versions of a 3D-printed stacking device called a ‘cassette’ were developed that are compatible with the variants of chips (Figure 3). The cassettes can be readily carried in 15 mL (96-well format chip) or 50 mL (24-well format chip) plastic tubes allowing a portable way to hold ten sandwiched chips for storage and transportation of crystals to the synchrotron at room-temperature. Lastly, to facilitate mounting of the chips onto a beamline goniometer for diffraction data collection, a holder was fabricated based on the previously reported goniometer-compatible flow-cell device (Ghosh *et al*., 2023). Briefly, the 3D-printed nozzle of the flow-cell was altered and equipped with grooves to hold the chips tailored for various chip-sizes and arrays of chips (Figure 2D). During data collection, a magnet is inserted at the base of the holder. The holder is mounted to the goniometer in the experimental hutch of the beamline and operated using standard software (Figure 2E, F).

**Figure 1.**
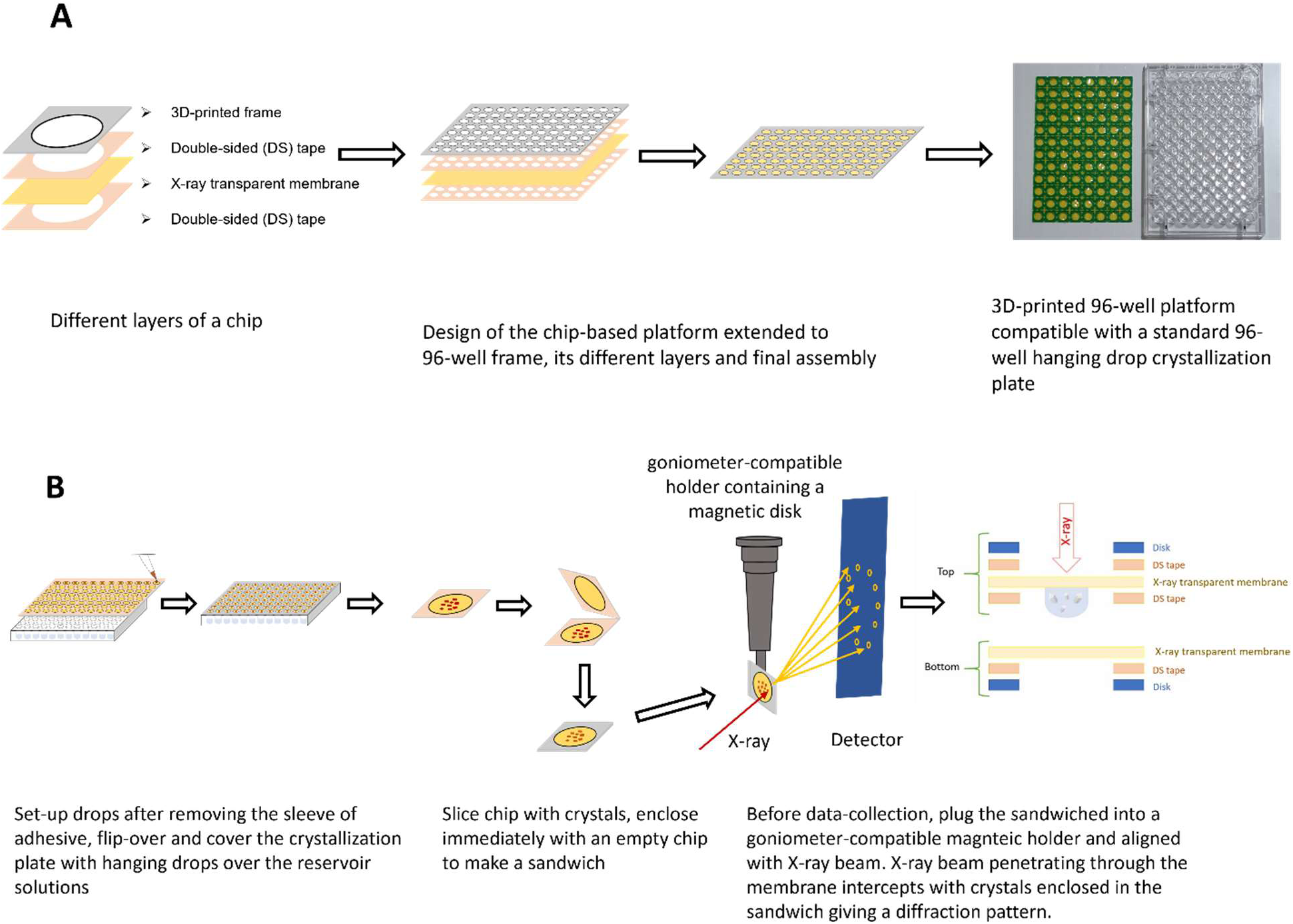
Design and use of the fixed-target device. **(A)** Design, development and fabrication of the frame consisting of a 3D-printed frame (with disk), that is assembled with double-sided (DS) tapes, and an X-ray transparent membrane. Once all the parts are assembled, they form a frame that can be either used for direct crystallization screening on top of crystallization plates or can be attached directly to a bottom frame with the sample in the middle for data collection. **(B)** Illustration showing the use of the platform for high-throughput hanging drop crystallization, crystal storage, and sample-delivery to the X-ray beam using a goniometer-compatible holder for diffraction data collection. The design results in a pathlength of ∼75 micrometers of the X-ray beam through the crystals in the enclosed chip.

### 2.2 Preparation of protein crystal samples

#### Lysozyme

Lysozyme was crystallized as reported previously (Diamond, 1974). Lysozyme from hen egg-white was purchased from WAKO Chemical Corporation (Lot PDN2655) and dissolved in 0.05 M sodium acetate buffer (pH 4.5) to a final concentration of 80 mg/ml and filtered through a 0.22 μm filter to remove particulates. On a 96-well plate, 100 μL of the crystallization solution was pipetted into the reservoir wells to obtain a concentration of 0.1 M sodium acetate (pH 4.5) and 1-2.5 M NaCl. A 1.5 μL drop of 1:1 mixture of lysozyme and reservoir solution was placed on each of the chips, after which the chip was flipped and pressed over the well of the plate. Crystals of lysozyme were obtained at 40 mg/mL in 50 mM sodium acetate (pH 4.5); 0.8 M NaCl at 20 °C in a few hours that grew to 150–200 μm (Figure 4A) in size after overnight incubation.

#### Photosynthetic reaction center from *Blastochloris viridis* (RC*_vir_*)

Growth, purification and crystallization of RC*_vir_* are described previously (Dods *et al*., 2017). The *B.viridis* cells were grown in an anaerobic environment for 36 hours in the dark and 48 hours in the light. Cells were harvested, disrupted by sonication and centrifuged to isolate membranes containing RC*_vir_*. Membranes were solubilized overnight in Tris buffer containing lauryldimehtylamine-N-oxide (LDAO) detergent and centrifuged. The protein was purified using anion exchange chromatography and gel filtration. For crystallization, a protein concentration of 7.5 mg/mL protein was used together with a crystallization solution (400 μL heptane-triol, 20 μL 1M KPi pH 6.8 and 475 mg Ammonium Sulphate) and crystal seeds. Two methods of crystallization were utilized (Figure 4B). Firstly, RC*_vir_* crystals were grown in sitting drop at 4°C and 3-5 μL of crystal slurry was pipetted on the chip. Alternately, microcrystals were generated at 4°C directly onto the chip using the hanging drop method with additional 2M ammonium sulfate in the reservoir.

#### ba_3_-type cytochrome c oxidase Thermus thermophilus (Tt CcO)

Microcrystallization of *Tt* C*c*O in LCP is prepared by modifying the previously reported method (Andersson *et al*., 2017). Briefly, *Tt* C*c*O was extracted and purified from bacterial membranes using affinity purification with Ni– nitrilotriacetic acid followed by overnight dialysis at 4°C and ion-exchange chromatography using a HiPrep DEAE FF 16/10 column. Purified protein was concentrated to 12–15 mg/ml and crystallized in LCP using a well-based technique in glass plates at room temperature as described previously (Andersson *et al*., 2019). The crystallization buffer contained 0.1 M MES (pH 5.3), 1.4 M NaCl and 39-41 % v/v PEG 400. Increase in PEG 400 concentration led to melting of LCP worms and helped to form crystals of slightly bigger size with 35-40 μm in their longer dimension. The crystals from melted LCP worms were harvested after 2–3 days, packed into a PCR tube and carried to the synchrotron.

### 2.3 X-ray diffraction data collection, processing and structure determination

On-chip grown lysozyme crystals were sandwiched, mounted, and in the case of cryogenic data collection the chip was directly frozen in the cryo stream without addition of cryoprotectant. Data collection on single crystals were performed at the Photon Factory (PF) BL-5A beamline at cryogenic temperature (Hiraki *et al*., 2008) and the PF-Advanced Ring (AR) AR-NW12A beamline at room-temperature (Chavas *et al*., 2012) of Japan’s High Energy Research Organization (KEK) radiation facilities. The beamline control software UGUI is available at both beamlines for sample viewing, alignment and diffraction measurements. At BL-5A, an X-ray beam size of 100 µm (V) × 100 µm (H) at 1.0 Å wavelength, an exposure time of 0.1 second per frame, flux of 1 × 10^11^ photons/s and 100% transmission was used to collect a total of 900 images. The chips were rotated with 0.1 degrees of oscillation from 45-135 degrees and diffraction data were collected on a Dectris PILATUS3 S6M detector. For the room-temperature data collection at AR-NW12A, we used an X-ray beam of 200 µm (V) × 130 µm (H) at 0.75 Å wavelength, exposure of 0.1 second per frame, a flux of 5 × 10^11^ photons/s and 100% transmission for the collection of 800 frames. Diffraction data were collected on a PILATUS3 S2M detector while rotating the chips from 50-130 degrees with 0.1 degrees of oscillation. Datasets were auto-processed using scripts adapted within the in-line PREMO (PF REmote Monitoring ref?) system to generate MTZ files. MTZ files were truncated and molecular replacement with PDB model 3WUN using Phaser in CCP4i (version 8.0.010) was employed. The structures were refined in CCP4i as well as using the PHENIX suite (version 1.17.1-3660).

Serial synchrotron crystallography on *Tt* C*c*O and RC*_vir_* were performed at the BioMAX beamline of MAXIV laboratory (Sweden) (Ursby *et al*., 2020) and the ID29 beamline of ESRF (France) (de Sanctis *et al*., 2012, Orlans *et al*., 2025). 2 μL of microcrystals were sandwiched on the chips (Figure 4C) which were mounted on a holder and aligned with the X-ray for data collection at room-temperature. Crystals were viewed on MxCuBE3 (Mueller, 2017) and appropriately sized grids were selected for data collection. At BioMAX, data collection was performed with X-ray beam size of 20 µm (V) × 20 µm (H), wavelength 0.98 Å (or photon energy of 12.7 keV) and a flux of 3.81 - 3.88 × 10^12^ photons/s, 100 % transmission, and recorded on an EIGER 16M hybrid pixel detector at a frame rate of 11 ms per frame. At ESRF, an X-ray beam size of 4 µm (V) × 2 µm (H), a photon energy of 11.56 keV, a flux of 2 × 10^15^ photons/s, a frequency of 231.25 Hz with a pulsed beam of 90 ms and a Jungfrau 4M detector was used. CrystFEL version 9.0 and above (White *et al*., 2012) integrated within the pipelines of the beamlines was used for indexing, integration, merging and conversion to MTZ format. MTZ files were truncated, the structures solved by molecular replacement using Phaser and refined with the CCP4i (Agirre *et al*., 2023) and CCP4 cloud (Krissinel *et al*., 2022) suite. Previously known crystal structures with PDB IDs 5NJ4 and 5NDC were used as models of RC*_vir_* and *Tt* C*c*O, respectively.

For all datasets, model building was performed in COOT (Emsley & Cowtan, 2004) and structural illustrations were drawn in PyMOL (DeLano, 2002). Final statistics of all the datasets are presented in Table 2.

## 3. Results and discussion

Throughout this work, we aimed to simplify the steps required for SX data collection at synchrotron facilities by the development of a fixed-target platform that is pre-assembled, convenient to use, can facilitate shipment of samples at room temperature, and can be mounted upon a standard SPINE magnet for data collection. In this manner, any X-ray diffraction beamline with a rapid readout X-ray detector could be adapted for SX data collection by making use of the raster-scanning capability of the goniometer. We demonstrate the versatility of this chip-based platform by applying various data-collection strategies at several synchrotron radiation sources targeting different proteins.

### 3.1 Assembly of the chip and hanging-drop crystallization

To minimize crystal handling, we developed a platform that could support hanging drop crystallization experiments above a crystallization well of a standard crystallization plate, and where the same frame could be used as the mount for X-ray diffraction data collection. We first explored this concept using a 24-well format (Figure 2A) and then extended the concept to the 96-well format (Greiner and TPP) plate for hanging drop crystallization setups.

For the 96-well design, a frame consisting of an array of flat circular supports (Figure 2B) was assembled with a layer of X-ray transparent membrane (Mylar or Kapton) of variable thickness (3-12 µm) using double-sided adhesive tape. The thickness of the 3D-printed support (∼0.75 mm) was chosen for its light weight and flexibility and could be easily detached with a scalpel for assembly of the platform. Our assembly has two types of frames: one top frame with a ‘support-adhesive-membrane-adhesive’ design that is suitable for on-chip crystallization, while the bottom frame has only one layer of double-sided tape (‘support-adhesive-membrane’) to be used for encapsulating pre-grown crystals within a sandwich to prevent the crystals from drying out. The adhesive layer within the sandwich provides a spacer of 75 µm for the encapsulated crystals. Two top frames can be used to create a chip for data collection that provides extra space for the sample if needed and was used in this study for the high-viscosity *Tt* C*c*O sample crystallized in LCP.

The 3D printed backbone support with adhesive fits onto a 96-well plates and the double-sided tape successfully creates an environment for vapor diffusion. This design thus supports *in situ* crystallization and the double-sided tape ensures a closed environment, with no grease being required to seal the chambers. The X-ray transparent membrane (3.0 to 7.6 µm thick Mylar or 12.6 µm thick Kapton) maintains a humid environment for a crystallization drop of ∼1.5 μL when using a 100 μL reservoir, and there was no indication of drops drying out after three weeks upon visual inspection. Crystal nucleation and growth can be directly monitored on the chips using a light microscope since Mylar is transparent to visible light. Once crystals appear, the chip is detached from the crystallization plate using a scalpel and sandwiched with a second chip to encapsulate crystals between the two membrane layers, which also prevents the samples from drying out after assembly (Figure 2C). This supported sandwich is then clipped onto a holder that is compatible with a SPINE base (Figure 2D) and data are collected using a 2D raster scan or by rotation of the chip (Figure 2E, F).

**Figure 2.**
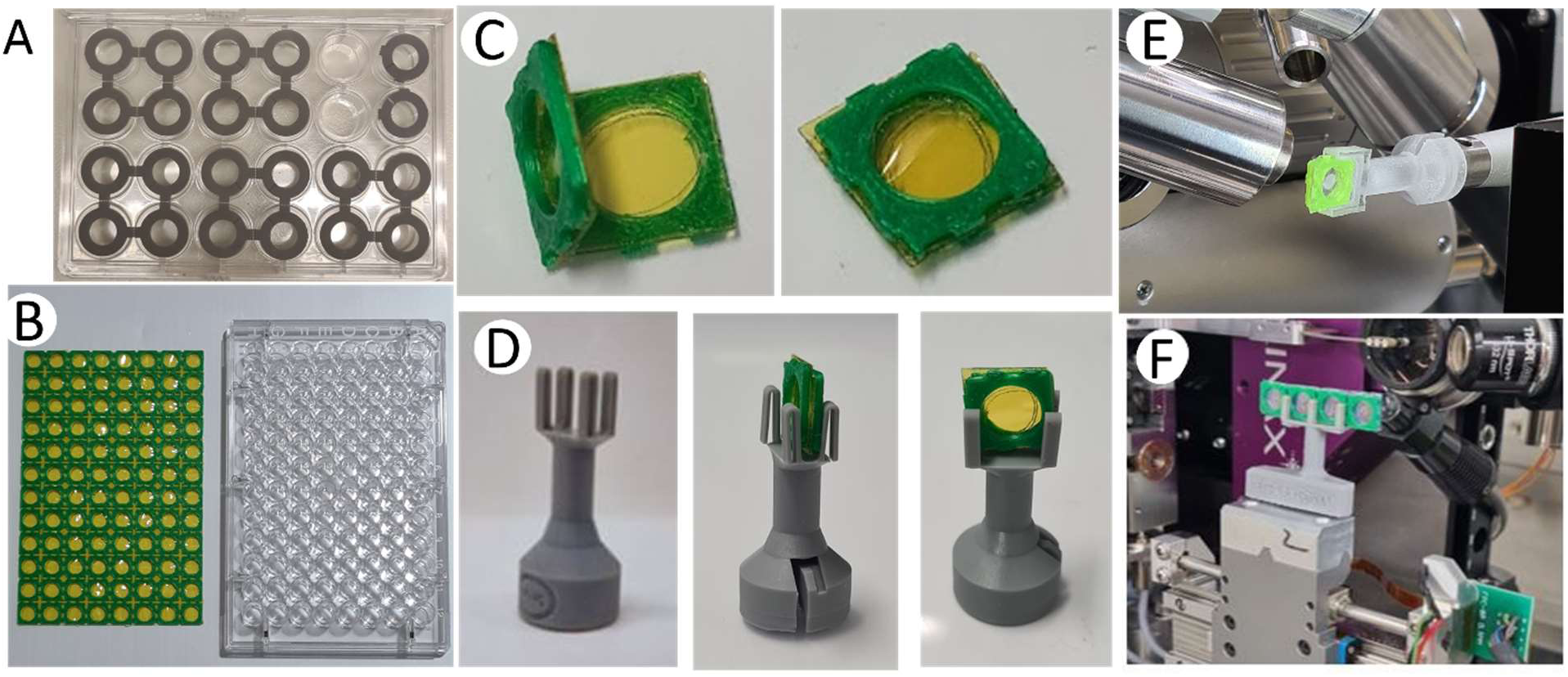
Design and assembly of the chip. **(A)** The 24-well design of the chip. **(B)** The 96-well design with a top frame next to a crystallization plate. **(C)** Chip assembly. **(D)** The chip holder with and without the chip inserted. **(E)** The chip mounted on the goniometer. **(F)** Multiple chips mounted on the goniometer.

The design is also available in a 24-well format. However, the 96-well framework is advantageous for several reasons. As it matches standard 96-well crystallization plates (Greiner and TPP) it is compatible with multi-channel pipettes and sample-dispensing robotics, and the device could potentially be used for high-throughput crystallization screening. Moreover, the 3D-printed support contains perforations between the individual squares that allows the frame to be separated into different parts so that one frame can be used to screen several different proteins or crystallization conditions. During crystallization on the 96-well frame, it is suggested to use a humidifier for sensitive proteins or buffers containing volatile additives to avoid drying of sample during crystallization.

A single crystallization drop can be extracted from the crystallization setup by cutting out the 3D-printed plastic frame of a single well. The single frame can then be sealed with a complementary frame, thus sandwiching the drop, to form a chip ready for data collection. Alternatively, it is possible to remove the entire 96-well frame from the crystallization plate and seal it with a complementary frame. Another variant is to leave every second well empty during crystallization, to be able to cut out two adjacent wells at a time and fold them onto each other so as to sandwich the crystallization drop (Figure 2C).

Importantly, the platform also allows the mounting crystals that have been grown using other methods (Uwangue *et al*., 2025). In this case, typically ∼1.5 µL of microcrystals are harvested from a crystallization plate or tube and pipetted directly onto the membrane of the support. In the case of LCP-grown crystals, they are extruded onto the membrane from a Hamilton syringe. An adjacent single frame can then be folded onto the crystal drop to seal it closed. Alternatively, all 96 wells are loaded with crystals and another 96-well support is used to sandwich all 96 drops in one step. In the latter case, once sealed, the sandwiches can be cut out individually and clipped on a holder, which in turn is mounted on the goniometer (Figure 2D and E). While the mounting may initially seem to involve multiple steps, the procedure requires minimal user training and significantly reduces the risks of crystals being lost or becoming dehydrated during assembly or data collection. To assemble the chips, the only tool that is required is a scalpel to cut out the frames.

### 3.2 Crystal storage and transportation

To store and transport the crystal-containing sandwiched chips at room-temperature, we constructed a 3D-printed compact device consisting of a stack of chip-holders or ‘cassettes’. This cassette is fitted inside a 15 mL Falcon plastic tube (Figure 3A, B) and a cotton pad soaked with crystallization solution or buffer is added to the tube to retain humidity around the crystals during storage and transportation, and the tube is sealed tightly. Lysozyme crystals shipped in the storage device did not dry out during shipment based on visual inspection (Figure 3C), and transported crystals of *in situ* grown RC*_vir_* retained diffraction quality after transportation in the storage system.

**Figure 3.**
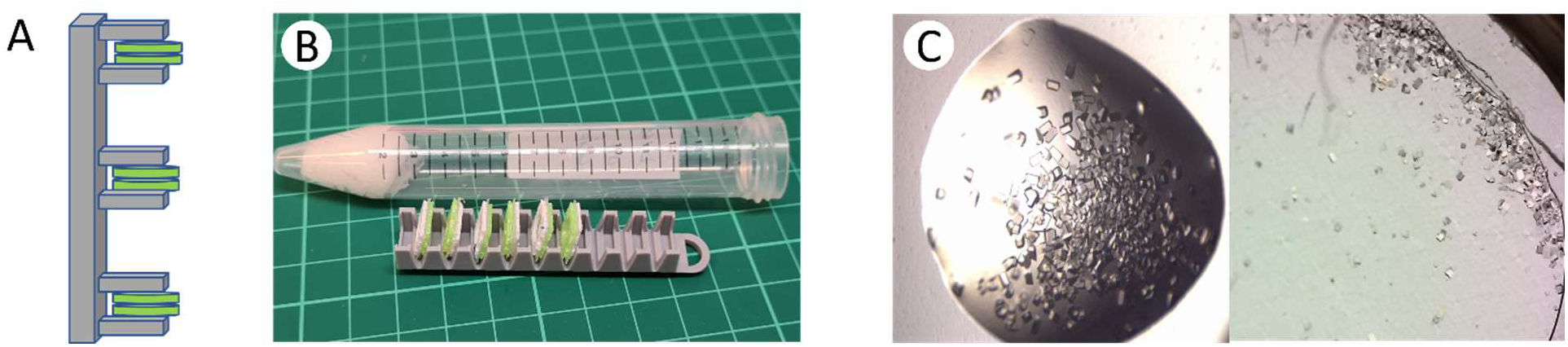
Shipment and storage of crystal-containing chips. **(A)** Cassette (grey) for storage and transportation of the assembled sandwiched chips (green). **(B)** Once the chips are clipped onto the holder, the entity is into a plastic tube containing a moist cotton/paper soaked with crystallization solution to prevent the samples from drying out. A 3D-printed holder for the 96-well sandwiched chips compatible with a 15 mL plastic tube is shown. **(C)** Crystals of lysozyme grown on the chip were captured before (left) and after (right) shipment of the sandwiched chips from the home laboratory (Gothenburg, Sweden) to the ID29 beamline of ESRF (Grenoble, France) with a total duration of ∼7 days.

### 3.3 Mounting of the chip for data collection

To mount the chip on the goniometer of a standard macromolecular crystallography synchrotron beamline, we adapted a design of a goniometer-compatible flow-cell (Ghosh *et al*., 2023) to be able to hold the sandwiched chips for alignment and data collection (Figure 1B). More specifically, the 3D-printed part was modified to hold various chip-sizes and arrays of chips (Figure 2E and F). The height of the holder is compatible with the SPINE standard (Beteva *et al*., 2006) to prevent any perturbation of the beamline optics and the mounting system can easily be adapted to the specific needs of the beamline. For data collection, the sandwiched chips, either assembled at the beamline or transported in the cassette, are slotted into the groove of the holder and mounted on to the goniometer using a magnetic disk at the base of the holder. At some beamlines it is possible for the scan domain of the goniometer to mount an array of chips which allows a more efficient data collection (Figure 2F).

### 3.4 X-ray diffraction data collection and resulting structures

As part of this study, we tested the versality of our fixed-target platform for various X-ray diffraction data-collection strategies at room- and cryo-temperature and on crystals from three types of proteins out of which one is crystallized in LCP. The results are summarized in Table 2. The first example is with the model protein lysozyme (Figure 5A). Using rotational crystallography, we collected datasets at cryogenic- and room-temperature from on-chip grown single crystals of lysozyme prepared on-site one day before data collection at the Photon Factory (PF) synchrotron radiation facility in Japan. Each chip contained several crystals of 150 µm × 200 µm × 200 µm in size (Figure 4A), and one crystal for each temperature was selected for rotational data collection. For the cryogenic data collection, crystals mounted on the chip were frozen at 100 K using cold vapor from the cryo-stream generator without the addition of any cryo-protecting solution. Depending on the crystallization condition, crystals may need to be complemented with a suitable cryo-protectant before data collection at low temperature. During data collection of on-chip grown lysozyme crystals at room-temperature, we noticed slight movement of crystals enclosed within the sandwiched chip when mounted on the goniometer. This was resolved by removing the excess volume of crystallization solution using a thin capillary wire from the sandwiched chip prior to data collection. It is noteworthy that during data collection, an opaque edge near the frame of the chips prevented data collection from a complete 360-degree rotation of the chip. This problem could be overcome by collecting wedges of data from both sides of the chip, or alternatively from several different crystals on a chip. For lysozyme, an ∼80 degrees rotation of a single crystal was sufficient to obtain nearly complete data at both temperatures. The resulting lysozyme structures were refined to resolutions of 2.3 Å and 1.6 Å for cryo- and room-temperature, respectively. From these rotational crystallography results it is clear that our chip-based platform allows *in situ* data collection from on-chip grown crystals at both room- and cryo-temperature.

Crystals of RC*_vir_* grown directly on the membrane surface of the chip differ in morphology and size compared to crystals obtained by the sitting-drop crystallization method (Figure 4B). The crystals obtained through on-chip crystallization were larger in size, and a complete dataset was collected at the BioMAX beamline of MAXIV Laboratory, Sweden, resulting in a 2.8 Å structure (Figure 5B). For comparison, RC*_vir_* microcrystals grown using sitting-drop vapor diffusion were applied to either the 24-well plate frame for data collection at BioMAX or the 96 well-plate frame for data collection at the ID29 beamline of ESRF. In both cases, data collection was performed by raster scanning with a mesh chosen according to the maximum area that the frame allowed. Both RC*_vir_* structures solved from sitting-drop microcrystals were refined to a resolution of 3.3 Å.

LCP grown crystals of *Tt* C*c*O were pipetted onto the membrane and the high-viscosity sample was efficiently spread out over the surface by gently pressing the membranes together as the chip was sealed (Figure 4C). The frame is closed by sandwiching it with another, causing the crystals to spread out over the surface. The volume of LCP applied to the chip in this case was a couple of microliters but has to be adopted based on the viscosity of the sample. To obtain the maximum number of images from one single chip, several grids can be drawn to cover the surface as efficiently as possible. The *Tt* C*c*O data presented here were collected from three chips where one to three grids were raster scanned per chip. The resulting structure (Figure 5C) at 2.3 Å resolution agrees well with previously solved structures of *Tt* C*c*O solved using a capillary-based flow cell (Ghosh *et al*., 2023).

**Table 2.**
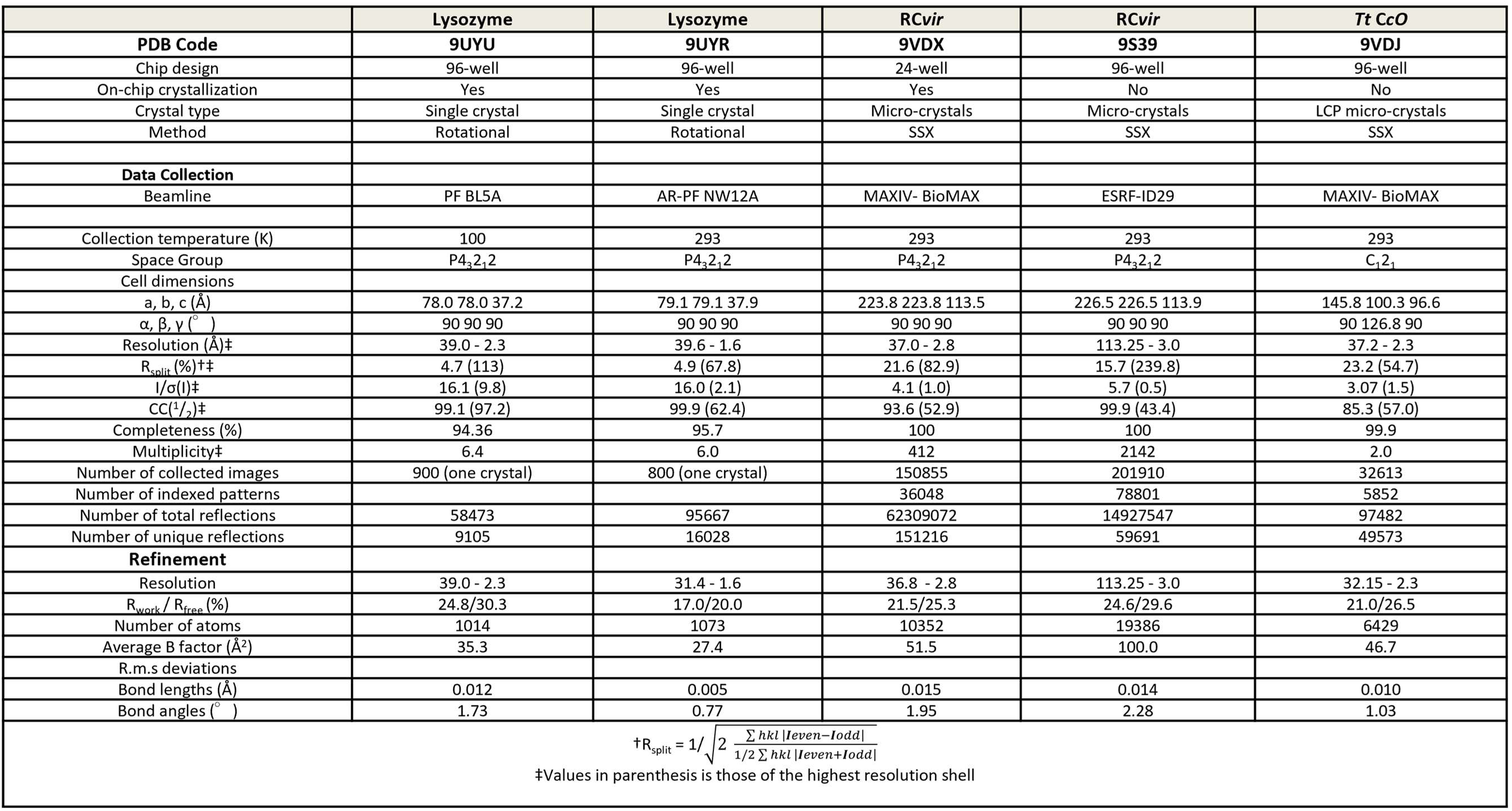
X-ray diffraction data collection and refinement statistics.

**Figure 4.**
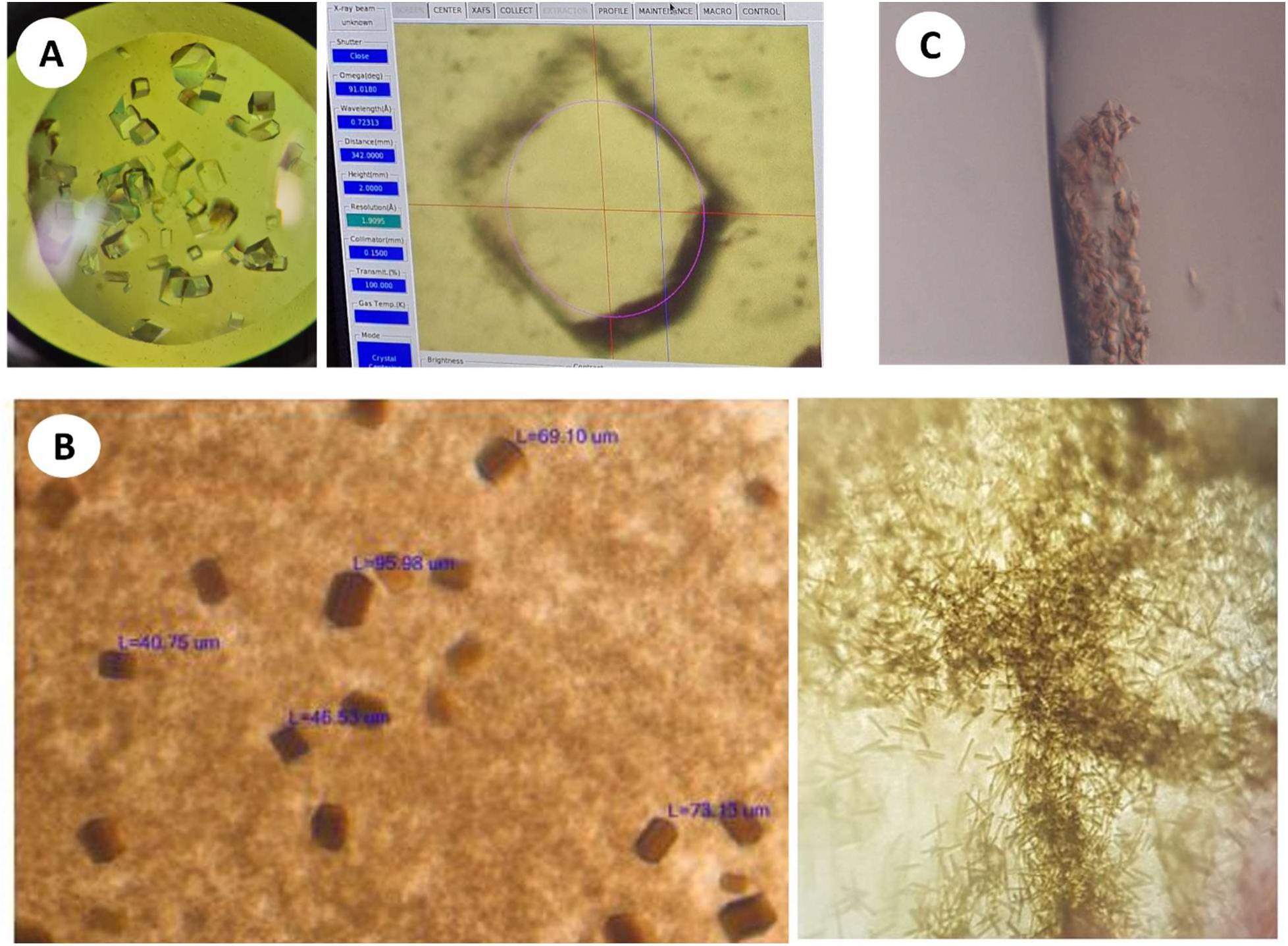
Crystals used for data collection. **(A)** Crystals of lysozyme of ∼150 × 200 × 200 µm^3^ size in a hanging drop visualized on a 96-well chip (left). A single crystal is selected for data collection at PF-AR NW12A beamline, Japan (right). **(B)** Crystals of RC*_vir_* approximately 20 × 40 × 80 µm^3^ in size were obtained from on-chip crystallization by hanging drop technique on a 24-well chip (left). Crystals of the same protein obtained in a sitting-drop plate with a size of ∼20 × 20 × 100 µm^3^ (right). **(C)** LCP crystals of *Tt* C*c*O visualized on the chip.

#### Comparison with other available chip options

A wide variety of alternatives are available for users interested in performing SX data collections using fixed-target devices, including some commercially accessible alternatives. In Table 1, a selection of these are listed in addition to the fixed-target chip presented in this work. The selected variants display differences regarding which data collection strategies they allow for, if they are compatible with on-chip crystallization or not and whether they are compatible also with the collection of cryo-temperature data. Out of the fourteen options presented, all but five require specialized hardware or stages to mount the sample onto the goniometer at the beamline. The other five options, including our device, instead take advantage of the SPINE system that is readily available at most macromolecular crystallography beamlines. This alternative makes the setup straightforward and more suitable also for non-expert users. A second measure of convenience is the possibility of crystallization directly on the device, *i.e. in situ* data collection without transfer of the crystals. In this case, four of the listed devices offer this possibility, including our chip. *In situ* data collection can save time and may aid in cases where the crystals are fragile and sensitive to handling. Another property that is listed is whether the device is disposable or not. There are pros and cons regarding using a disposable device. The advantage is that a disposable device minimizes the risk of cross-contamination between samples and eliminates the need for time-consuming cleaning procedures, improving efficiency. In contrast, single-use devices generate more waste. Most of the devices listed rely on the raster-scan data collection strategy, yet three of the devices presented including ours are also compatible with rotational data collection. Which of these strategies is most suitable depends on many factors such as the nature of the sample including crystal size and sample consistency. An issue that was observed in this study is that crystals contained in liquid on some occasions moved within the chip during data collection. This problem was to a large extent mitigated by use of the smaller-size 96-well instead of the 24-well chip and optimization of the crystal volume applied to each chip. Some of the devices shown in Table 1 resolved this issue by using a specialized loading system, although this might complicate the sample preparation process. Finally, it should be noted that the chip presented here, unlike most devices listed in Table 1, is compatible with LCP crystals and allowed a high-quality structure of the *Tt* C*c*O membrane protein to be solved.

## Conclusion

In this study, we present a fixed-target device developed for serial crystallography data collection at synchrotron radiation sources. The chip is easy to use, less fragile than many alternatives, allows on-chip crystallization and enables high-quality X-ray diffraction data to be collected by use of the SPINE-based system available at most relevant beamlines. Importantly, it enables diffraction data to be collected from very low amounts of crystal sample. The benefits of the chip are presented by showcasing results from three different protein systems including a membrane protein crystallized in LCP. Data were collected using different strategies at both cryo- and room-temperature to verify the versatility of the device. Finally, a simple system to transport the pre-loaded chips at room-temperature and with retained humidity is presented.

**Figure 5.**
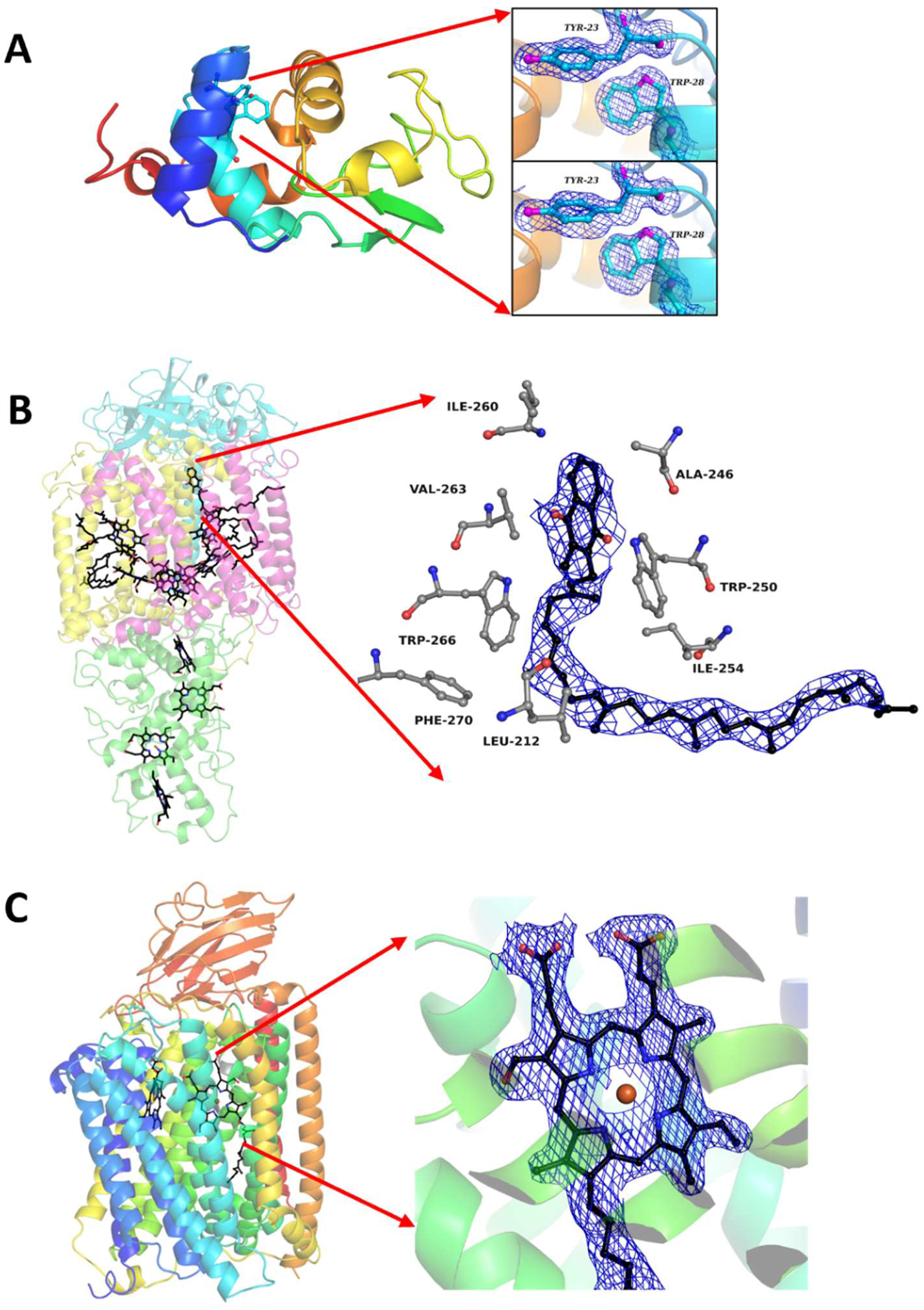
Structures of lysozyme, RC*_vir_* and *Tt* C*c*O. **(A)** Crystal structures of *in situ* grown lysozyme single-crystals at cryogenic (top) and room-temperatures (bottom). 2Fo-Fc (blue, 1.0 σ) electron density maps of selected residues are shown. Serial synchrotron crystallography structures of **(B)** RC*_vir_* and **(C)** *Tt* C*c*O with a co-factor highlighted in each case. The 2Fo-Fc electron density is contoured at 1.0 σ (blue).

## Acknowledgements

X-ray diffraction data were collected at the BioMAX beamline of MAX IV Laboratory (proposal number 20220186), ID29 of the ESRF (BAG proposal number MX-2543) and Photon Factory (PF) and PF-Advanced ring synchrotron beamlines BL-5A and AR-NW12A (proposal number 2022G575). We extend our gratitude to the staff of MAX IV, ESRF, PF and PF-AR for their support during experiments.

